# *In silico* and functional characterisation of an ultra-rare *CFTR* mutation identifies novel lasso motif interactions regulating channel gating

**DOI:** 10.1101/2021.12.12.472297

**Authors:** Sharon L. Wong, Nikhil T. Awatade, Miro A. Astore, Katelin M. Allan, Michael J. Carnell, Iveta Slapetova, Po-chia Chen, Alexander Capraro, Laura K. Fawcett, Renee M. Whan, Renate Griffith, Chee Y. Ooi, Serdar Kuyucak, Adam Jaffe, Shafagh A. Waters

## Abstract

Characterisation of I37R – a novel mutation in the lasso motif of ABC-transporter CFTR, a chloride channel – was conducted by theratyping using CFTR potentiators which increase channel gating activity and correctors which repair protein trafficking defects. I37R-CFTR function was characterised using intestinal current measurements (ICM) in rectal biopsies, forskolin-induced swelling (FIS) in intestinal organoids and short circuit current measurements (I_sc_) in organoid-derived monolayers from an individual with I37R/F508del CFTR genotype. We demonstrated that the I37R-CFTR mutation results in a residual function defect amenable to treatment with potentiators and type III, but not to type I, correctors. Molecular dynamics of I37R-CFTR using an extended model of the phosphorylated, ATP-bound human CFTR identified an altered lasso motif conformation which results in an unfavourable strengthening of the interactions between the lasso motif, the regulatory (R) domain and the transmembrane domain two (TMD2). In conclusion, structural and functional characterisation of the I37R-*CFTR* mutation increases understanding of CFTR channel regulation and provides a potential pathway to access CFTR modulator treatments for individuals with CF caused by ultra-rare *CFTR* mutations.

## Introduction

Cystic fibrosis (CF) is a life-limiting genetic disease resulting from mutations in the CF transmembrane conductance regulator (*CFTR*) gene (Ratjen et al. 2015). CFTR – the only member of the ABC transporter family known to be an ion channel – consists of two transmembrane domains (TMD1 and TMD2) which form an anion-selective pore, two highly conserved nucleotide-binding domains (NBD1 and NBD2) with ATP-binding pockets and a newly described N-terminal lasso motif (Hwang and Kirk 2013; Zhang and Chen 2016). In addition, CFTR has a unique, disordered regulatory (R) domain which contains protein kinase A (PKA) phosphorylation sites. For the CFTR channel to open and close (gate), cAMP-dependent PKA phosphorylation of the R domain first activates the CFTR (Gadsby and Nairn 1994). Then, ATP-binding induces the dimerization of the two NBDs which opens the channel pore and ATP hydrolysis closes the pore.

The lasso motif (amino acids (aa) M1-L69), which is partially embedded in the bilayer and interacts with the R domain, was recently resolved following advancements in cryo-electron microscopy (cryo-EM) of the CFTR structure (Liu et al. 2017; Zhang, Liu, and Chen 2018). The first 40 amino acids of the lasso motif, which include lasso helix 1 (Lh1, aa V11–R29), form a circular “noose” structure (Hoffmann et al. 2018). The noose structure wraps around the transmembrane helices (TM2, TM6 of TMD1 and TM10, TM11 of TMD2) and is held in place by hydrophobic interactions with L15, F16, F17, T20, L24 and Y28. The C-terminal end of the lasso, which includes the lasso helix 2 (Lh2, aa A46–L61), is tucked under the elbow helix (aa I70–R75) (Hoffmann et al. 2018). Variable disease severity and heterogeneous clinical presentation has been reported for the 78 *CFTR* variants identified so far in the lasso motif (CFTR1 and CFTR2 databases, **Supp material 1**). Evidently, the lasso motif has a multifunctional role in CFTR regulation with variants impacting folding, gating and stability of the CFTR protein (Jurkuvenaite et al. 2006; Sabusap et al. 2021; Fu et al. 2001; Gené et al. 2008; Thelin et al. 2007).

CFTR modulators, small molecules which directly target CFTR dysfunction, are now available to certain individuals with CF. Currently, two classes are approved; (1) potentiators, which open the channel pore such as ivacaftor (VX-770) and (2) correctors, which assist CFTR protein folding and delivery to the cell membrane. Type I correctors (lumacaftor/VX-809, tezacaftor/VX-661) stabilise the NBD1-TMD1 and/or NBD1-TMD2 interface by binding directly to TMD1 (Loo, Bartlett, and Clarke 2013; Ren et al. 2013) or NBD1 which improves the interaction between NBD1 and the intracellular loops (Loo and Clarke 2017; Hudson et al. 2017). Type II correctors (C4) stabilise NBD2 and its interface with other CFTR domains while type III correctors (elexacaftor/VX-445) directly stabilise NBD1 (Okiyoneda et al. 2013). Combination therapies of corrector(s) and a potentiator (Orkambi^®^, Symdeko/Symkevi^®^, Trikafta/Kaftrio^®^) have been approved for CF individuals with F508del, the most common *CFTR* mutation, as well as several specific residual function mutations. Most recently, Trikafta/Kaftrio^®^ has been approved for patients with a single F508del mutation in combination with a minimal function mutation, broadening the population of CF patients eligible for treatment with CFTR modulator therapy.

Mounting evidence has shown that *in vitro* functional studies in patient-derived cell models successfully predict clinical benefit of available CFTR modulators for individuals bearing ultra-rare mutations (Ramalho et al. 2021; McCarthy et al. 2018; Berkers et al. 2019). In individuals with CF, adult stem cells are usually collected by taking either airway brushings or rectal biopsies. Single Lgr5^+^ stem cells, derived from crypts within a patient’s intestinal epithelium, can be expanded in culture medium and differentiated into organised multicellular structures complete with the donor patient’s genetic mutation(s), thus representing the individual patient (Sato et al. 2009). Stem cell models can be used for personalised drug screening to theratype and characterise rare CFTR mutations (Pollard and Pollard 2018; Awatade et al. 2018; Berkers et al. 2019). Determining the functional response of rare, uncharacterised CFTR mutations to modulator agents with known CFTR correction mechanisms enables characterisation of CFTR structural defects and enhances our understanding of CFTR function.

I37R-*CFTR* is a novel missense mutation in the lasso motif, detected in an Australian male child diagnosed through newborn screening with elevated immunoreactive trypsinogen, raised sweat chloride (>60 mmol/L) and CFTR Sanger sequencing identifying c.1521-1523del (F508del) and c.110C>T (I37R) mutations (**Supp material 2**). We used functional studies and molecular dynamics (MD) simulations to characterise the functional and structural defects of I37R-CFTR. CFTR function was assessed using intestinal current measurements (ICM) in rectal biopsies, forskolin-induced swelling (FIS) assays in intestinal organoids and short circuit current measurements (I_sc_) in I37R/F508del organoid-derived monolayers respectively. The potentiators VX-770 (approved), GLPG1837 (phase II clinical trials) and genistein (a natural food component with potentiator activity (Dey, Shah, and Bradbury 2016)) were tested as monotherapies, dual potentiator therapies or in combination with correctors (VX-809, VX-661 and VX-445). We compared this to our laboratory reference intestinal organoids. For MD simulations, we modelled and examined the structural defect of the I37R mutation on an extended cryo-EM structure of ATP-bound, phosphorylated human CFTR (PDB ID code 6MSM) (Zhang, Liu, and Chen 2018).

## Results

### I37R-CFTR baseline activity in patient-derived rectal biopsies and intestinal organoids

Intestinal current measurements (ICM) were performed on I37R/F508del and reference CF (F508del/F508del, G551D/F508del) and non-CF (wild-type: WT/WT) rectal biopsies using a standard protocol (Clancy et al. 2013; Graeber et al. 2015) (**Fig 1A**). Following stimulation with a forskolin (fsk) and IBMX cocktail, rectal biopsies from the I37R/F508del CF participant elicited cAMP-dependent currents of 45.8 ± 3.8 µA/cm^2^ – an appreciable 50% of WT-CFTR activity (P<0.05; **Fig 1A, Supp material 3**). This response was at least 4-fold higher than those of the reference CF biopsies, although statistical significance was not reached.

**Figure 1.**
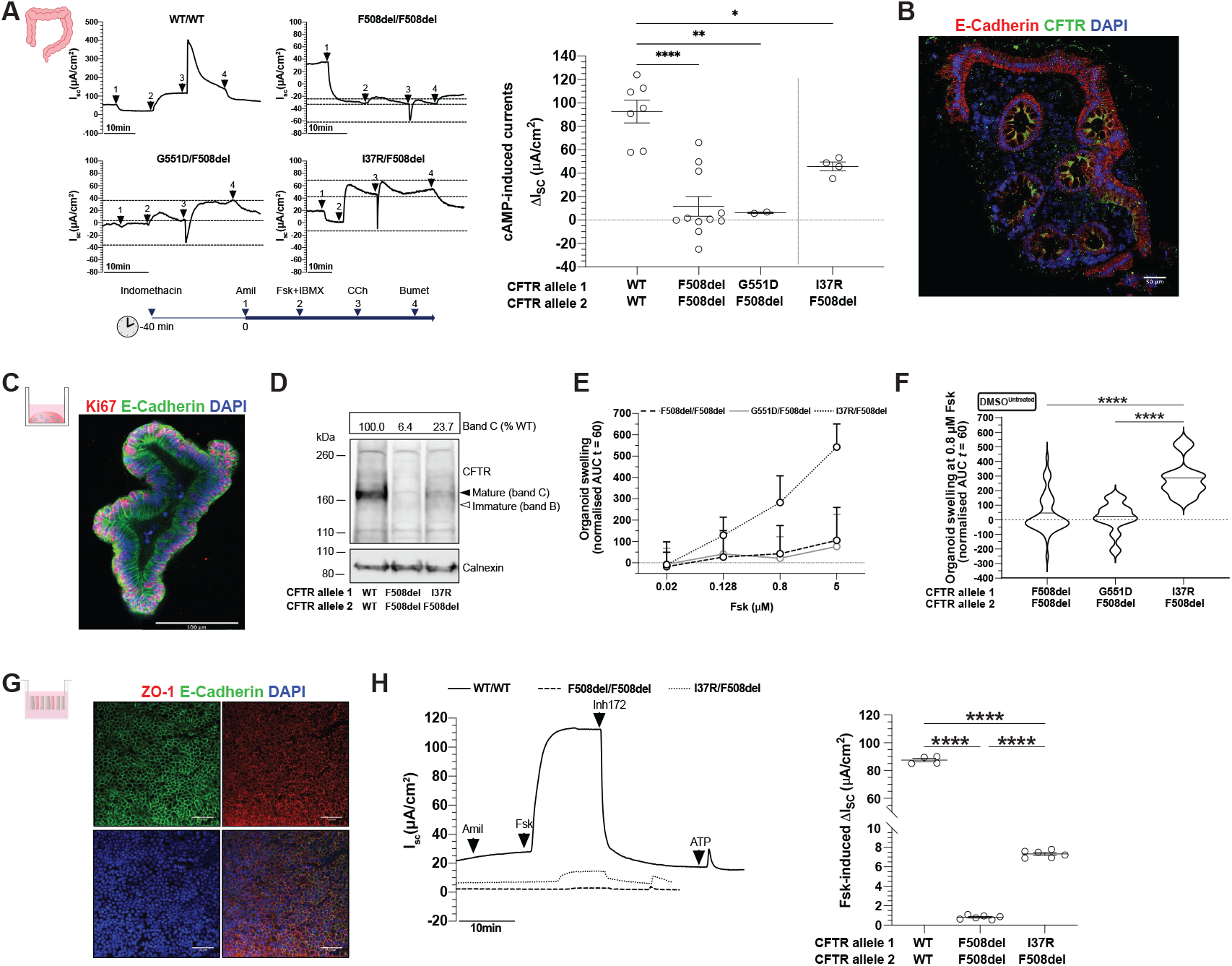
Characterisation of I37R-CFTR residual function in rectal biopsies and intestinal organoids. **(A)** Representative Ussing chamber recordings of intestinal current measurements (ICM) in rectal biopsies from WT-CFTR control participants and participants with CF. Dot plots of cAMP-induced current (ΔIsc-Fsk+IBMX) in participants with WT/WT (n=2), F508del/F508del (n=3), G551D/F508del (n=1) and I37R/F508del (n=1) CFTR genotypes. Experiments were performed in the presence of 10 µM indomethacin. Arrows indicate the addition of compounds: 100 µM apical amiloride (1. Amil), apical and basal addition of 10 µM forskolin + 100 µM IBMX cocktail (2. Fsk+IBMX), 100 µM basal carbachol (3. CCh) and 100 µM basal bumetanide (4. Bumet). The Isc at the time CCh was added (middle horizontal dotted line), and the maximum (top dotted lines) and minimum (bottom dotted lines) Isc induced are indicated. Each dot represents an individual replicate. **(B)** Immunofluorescence staining of CFTR (green), e-cadherin (red) and DAPI (blue) in a rectal biopsy derived from a I37R/F508del participant. 63x/1.4 oil immersion objective. Scale bar=50 µm. **(C)** Immunofluorescence staining of e-cadherin (green), Ki67 (red) and DAPI (blue) in intestinal organoids derived from a I37R/F508del participant. 20x/0.75 dry objective. Scale bar=100 µm. **(D)** Western blot in WT/WT, F508del/F508del and I37R/F508del intestinal organoids. CFTR maturation was calculated by measuring the level of mature mutant CFTR (Band C) as a percentage of mature CFTR from WT organoids (% normal CFTR). All data were normalised to the calnexin loading control. Band C represents the mature, complex-glycosylated CFTR. Band B represents the immature, core-glycosylated CFTR. **(E-F)** Forskolin-induced swelling (FIS) assay in organoids from participants with F508del/F508del (n=5), G551D/F508del (n=2) and I37R/F508del (n=1) CFTR genotypes. Organoids were stimulated with forskolin (fsk) concentrations ranging from 0.02 to 5 µM. **(E)** FIS expressed as the means ± standard deviation (SD) of the area under the curve (AUC) calculated from t=0 (baseline) to t=60. (F) FIS of organoids at 0.8µM fsk at baseline represent residual CFTR function. **(G)** Immunofluorescence staining of e-cadherin (green), ZO-1 (red) and DAPI (blue) in organoid-derived monolayers from a CF participant. 20x/0.75 dry objective. Scale bars = 50 µm. **(H)** Representative Ussing chamber recordings of short circuit current in organoid-derived monolayers from a WT-CFTR control participant and participants with CF. Dot plots of fsk-induced current (ΔIsc-Fsk) in participants with WT/WT (n=1), F508del/F508del (n=1) and I37R/F508del (n=1) CFTR genotypes. Experiments were performed in the presence of 10 µM indomethacin. Arrows indicate the addition of compounds: 100 µM apical amiloride, 5 µM basal fsk, 30 µM apical CFTR inhibitor CFTRinh-172 and 100 µM apical ATP. Each dot represents an individual replicate. One-way analysis of variance (ANOVA) was used to determine statistical differences. *P < 0.05, **P < 0.01, ****P < 0.0001.

Co-activation with carbachol (CCh) resulted in a biphasic response in the I37R/F508del biopsies, characteristic of residual CFTR chloride channel function in the CF colon (Veeze et al. 1994; Graeber et al. 2015). The initial negative I_sc_ peak indicates apical potassium secretion reached 9.4±2.5 µA/cm^2^. Following this, the CCh-induced positive I_sc_ indicates the increase of apical chloride secretion reached 15.78±2.07 µA/cm^2^. This biphasic response was similarly observed in the G551D/F508del biopsies (25.77±2.16 µA/cm^2^) but was diminished in the F508del/F508del biopsies (-2.28±1.65 µA/cm^2^). These findings are in accordance with the localisation of CFTR protein at the plasma membrane (mature complex-glycosylated CFTR) of the I37R/F508del rectal biopsies, as demonstrated by immunofluorescence staining (green; **Fig 1B**).

Next, CFTR protein expression and maturation was assessed in I37R/F508del, reference F508del/F508del and WT/WT organoids using western blot (**Fig 1C-D**). The expression of complex-glycosylated C band in I37R/F508del organoids was 23.7% that of the WT/WT organoids, considerably higher than the 6.4% detected from F508del/F508del organoids (**Fig 1D**). CFTR activity was then evaluated in I37R/F508del and CF reference intestinal organoids using a fsk-induced swelling (FIS) assay at four fsk concentrations between 0.02 to 5 µM (**Fig 1E**). FIS of I37R/F508del intestinal organoids at 0.8 µM fsk – the optimal concentration for baseline assessment of CFTR activity (Dekkers, Berkers, et al. 2016) – was 282.9±36.0 (**Fig 1E-F**). This exceeded the baseline FIS of the reference intestinal organoids by at least 7-fold (F508del/F508del: AUC=42.8±19.4; G551D/F508del: AUC=21.3±29.4).

The morphological difference between WT (pre-swollen) and CF organoids (Cuyx et al. 2021), means comparing CFTR activity between CF and healthy CFTR function by FIS assay cannot be achieved (van Mourik, Beekman, and van der Ent 2019; Dekkers, Berkers, et al. 2016). In order to compare I37R/F508del to wild-type CFTR activity, organoid derived monolayers were created (**Fig 1G)** and CFTR ion transport was performed (Zomer-van Ommen et al. 2018). Fsk-stimulated CFTR-dependent currents were 9-fold higher in I37R/F508del monolayers than those of reference F508del/F508del monolayers (7.3±0.2 vs 0.8±0.1 µA/cm^2^; P<0.0001), but 12-fold lower than WT/WT monolayers (87.5±1.3 µA/cm^2^; P<0.0001) (**Fig 1H**). This is consistent with the FIS assay results demonstrating high baseline CFTR activity in I37R/F508del intestinal organoids.

### I37R-CFTR functional response to CFTR modulator monotherapy in intestinal organoids

We investigated the functional response of I37R/F508del organoids to single potentiators – VX-770, GLPG1837 (G1837) and genistein (Gen). Treatment with VX-770 minimally increased FIS of I37R/F508del organoids by AUC of 59.7 above baseline at 0.128 µM fsk (**Fig 2A-C; Supp material 4)** – the optimal concentration for *in vitro* assessment of CFTR modulator response to predict clinical effect (Dekkers, Berkers, et al. 2016). G1837 and Gen both significantly increased FIS, albeit with different efficacies (655.8 and 256.8, respectively; **Fig 2A-C; Supp material 4)**. None of the potentiator treatments increased FIS in F508del/F508del organoids, indicating no improvement in CFTR activity in response to potentiator therapy (**Fig 2C**). Only G1837 significantly increased FIS in the G551D/F508del organoids (210.4±57.5; P<0.01). In comparison to G551D/F508del organoids, G1837 was 3-fold more efficacious in the I37R/F508del organoids (P<0.0001).

**Figure 2.**
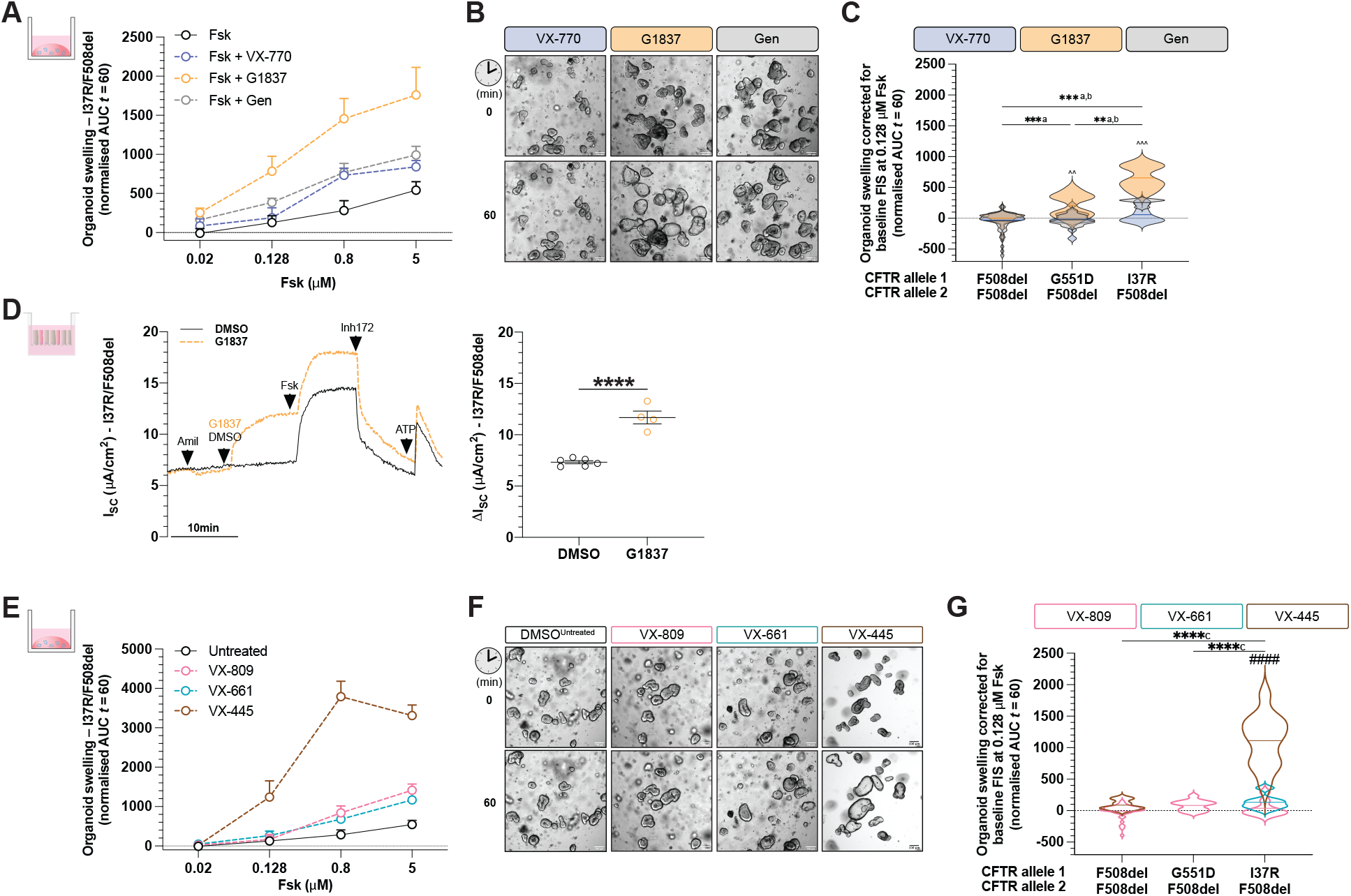
Characterisation of I37R-CFTR functional response to corrector or potentiator monotherapy in intestinal organoids. Forskolin-induced swelling (FIS) assay in organoids from participants with F508del/F508del (n=5), G551D/F508del (n=2) and I37R/F508del (n=1) CFTR genotypes. Organoids were incubated overnight with 0.03% DMSO (untreated) or 3 µM VX-809 or 3 µM VX-661 or 3 µM VX-445. After 24h, organoids were stimulated with fsk concentrations ranging from 0.02 to 5 µM, either alone or in combination with potentiator monotherapy (3 µM VX-770 or 3 µM G1837 or 50 µM Gen). **(A)** FIS of I37R/F508del organoids stimulated with VX-770, GLPG1837 (G1837) or genistein (Gen) monotherapy, expressed as the means ± standard deviation (SD) of the area under the curve (AUC) calculated from t=0 (baseline) to t=60 min. **(B)** Representative brightfield images of I37R/F508del organoids at baseline (t=0) and after 1h of stimulation (t=60) at 0.128 µM fsk. Scale bars=100 µm. **(C)** FIS of organoids at 0.128uM fsk following stimulation with VX-770, GLPG1837 (G1837) or genistein (Gen) monotherapy. Data corrected for baseline FIS and represented as violin plots to show distribution. **(D)** Representative Ussing chamber recordings of short circuit current in I37R/F508del organoid-derived monolayers. Dot plots of total currents stimulated by DMSO or G1837 plus fsk. Experiments were performed in the presence of 10 µM indomethacin. Arrows indicate the addition of compounds: 100 µM apical amiloride, apical addition of either vehicle control 0.01% DMSO or 10 µM G1837, 5 µM basal fsk, 30 µM apical CFTR inhibitor CFTRinh-172 and 100 µM apical ATP. Each dot represents an individual replicate. **(E)** FIS of I37R/F508del organoids pre-incubated with corrector (VX-809 or VX-661 or VX-445) for 24 h, expressed as the means ± standard deviation (SD) of the area under the curve (AUC) calculated from t=0 (baseline) to t=60 min. **(F)** Representative brightfield images of I37R/F508del organoids at baseline (t=0) and after 1h of stimulation (t=60) at 0.128 µM fsk. Scale bars=100 µm. **(G)** FIS of organoids at 0.128uM fsk following incubation with corrector (VX-809 or VX-661 or VX-445) for 24 h. Data corrected for baseline FIS and represented as violin plots to show distribution. One-way analysis of variance (ANOVA) was used to determine statistical differences. **P < 0.01, ***P < 0.001 and ****P < 0.0001. aP for G1837, bP for Gen and cP for VX-445 of I37R/F508del, ^P for G1837 vs VX-770 or Gen and #P for VX-445 vs VX-809 or VX-661.

Since G1837 demonstrated the greatest restoration of CFTR activity in I37R/F508del organoids, we evaluated G1837 treatment of I37R/F508del organoid-derived monolayers. G1837 led to a significant 1.5-fold increase in fsk-stimulated currents (ΔI_sc_: 4.4 µA/cm^2^; P<0.0001) (**Fig 2D**). This is consistent with the FIS of I37R/F508del organoids, indicating that I37R-CFTR responds to potentiator agents.

Given the I37R/F508del high residual CFTR activity and its localisation at the epithelial cell surface, we hypothesised that the I37R-*CFTR* mutation has minimal impact on CFTR protein folding or maturation. Treatment of I37R/F508del organoids with type I corrector agents (VX-809 or VX-661) did not significantly increase FIS above baseline (**Fig 2E-G; Supp material 4**). In contrast, treatment of I37R/F508del organoids with a type III corrector agent (VX-445) significantly increased FIS by AUC of 1112.5 above baseline, greater than those in the F508del/F508del organoids (42.5). VX-445 has been shown to act as both a corrector and potentiator for certain *CFTR* mutations (Shaughnessy, Zeitlin, and Bratcher 2021; Laselva et al. 2021; Veit, Vaccarin, and Lukacs 2021). Acute treatment of I37R/F508del organoids with VX-445 did not improve potentiation of CFTR (**Supp material 4**). This supports the observation that VX-445-stimulated rescue of CFTR in I37R/F508del organoids acts by a correction mechanism improving I37R mild folding and processing defects.

### I37R-CFTR functional response to CFTR modulator co-therapies in intestinal organoids

Combination treatments of CFTR modulators are used to treat patients bearing *CFTR* mutations with multiple functional defects such as F508del and patients who are heterozygous for CFTR mutations. We investigated the effect of combinations of potentiators. Dual potentiator combinations increased FIS of I37R/F508del organoids to a greater extent than the respective single potentiators (**Fig 3A**) and had a synergistic effect, where the FIS was greater than the sum of the respective single potentiators (**Supp material 5**). Despite G1837+Gen having greater efficacy than the other dual potentiator combinations, the magnitude of response was not statistically different between the different combinations of dual potentiators (**Fig 3A**).

**Figure 3.**
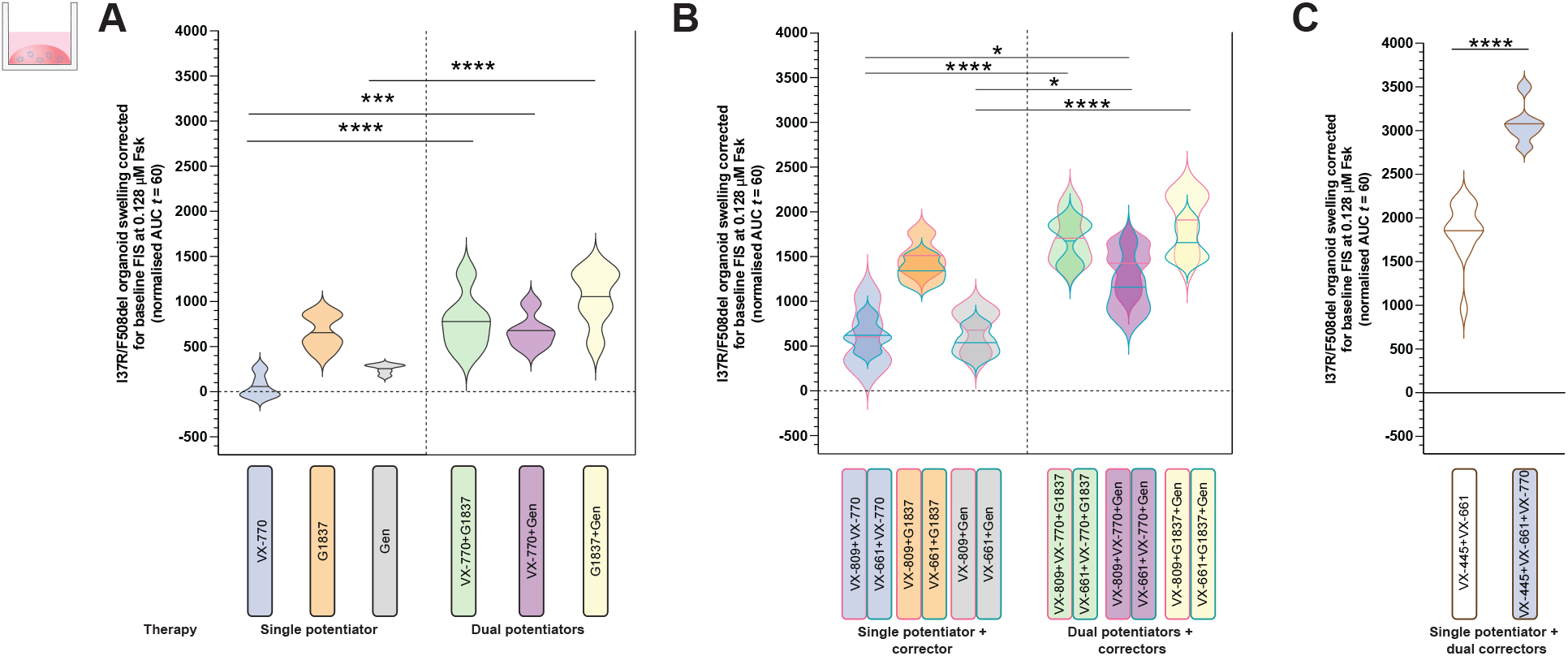
Characterisation of I37R-CFTR functional response to dual potentiator or corrector therapy or corrector(s)-potentiator(s) co-therapy in intestinal organoids. Forskolin-induced swelling (FIS) assay in organoids from participants with F508del/F508del (n=5), G551D/F508del (n=2) and I37R/F508del (n=1) CFTR genotypes. Organoids were incubated overnight with 0.03% DMSO (untreated) or 3 µM VX-809 or 3 µM VX-661 or 3 µM VX-445+18 µM VX-661. After 24h, organoids were stimulated with fsk ranging in concentration from 0.02 to 5 µM, either alone or in combination with a single potentiator (3 µM VX-770 or 3 µM G1837 or 50 µM Gen) or dual potentiators (VX-770+G1837 or VX-770+Gen or G1837+Gen). FIS of organoids at 0.128uM fsk stimulated with VX-770, GLPG1837 (G1837) or genistein (Gen) or their combinations, following **(A)** 24 h pre-incubation with DMSO (untreated) or **(B)** corrector (VX-809 or VX-661) respectively. **(C)** FIS of organoids at 0.128uM fsk stimulated without or with VX-770, following 24 h pre-incubation with dual correctors (VX-445+VX-661). Data corrected for baseline FIS and represented as violin plots to show distribution. One-way analysis of variance (ANOVA) was used to determine statistical differences. *P < 0.05, ***P < 0.001 ****P < 0.0001

Co-therapy with a corrector (VX-809 or VX-661) and dual potentiators significantly (P<0.01) increased FIS of I37R/F508del organoids compared to co-therapy of a corrector with VX-770 or Gen, but not G1837 (**Fig 3B**). VX-809/G1837+Gen co-therapy had the greatest efficacy, increasing FIS 1904.0 above baseline. In contrast, corrector/VX-770+Gen co-therapy had the least efficacy. This trend was consistent with that of the dual potentiators synergistic effect.

Dual correctors (VX-445+VX-661) increased FIS in I37R/F508del organoids by AUC of 1856.6 above baseline, which corresponds with the level of rescue achieved by the most effective corrector/dual potentiator co-therapy (VX-809/G1837+Gen). The triple combination therapy with dual correctors and a potentiator further increased FIS in I37R/F508del organoids by AUC of 3101.6 above baseline. It is therefore the most effective modulator combination tested in this study.

### I37R-CFTR perturbs the noose structure of the lasso motif

We next characterised the structural defect of I37R-CFTR using MD simulations. The primary structure of the lasso motif (M1-L69) is conserved across 230 vertebrate species (**Supp material 6-7**). The lasso motif formed a noose structure that rested against TMD2 (**Fig 4A**). Amino acids V12-R29 were embedded in the plasma membrane while the rest of the lasso motif resided in the cytosol. The noose structure was maintained by a salt bridge formed between K26 and D36 **(Fig 4B)**. I37 was positioned in the centre of this noose, within a hydrophobic pocket formed by amino acids from the lasso, TMD2, and the poorly resolved R domain in the cytosol (**Fig 4C**).

**Figure 4.**
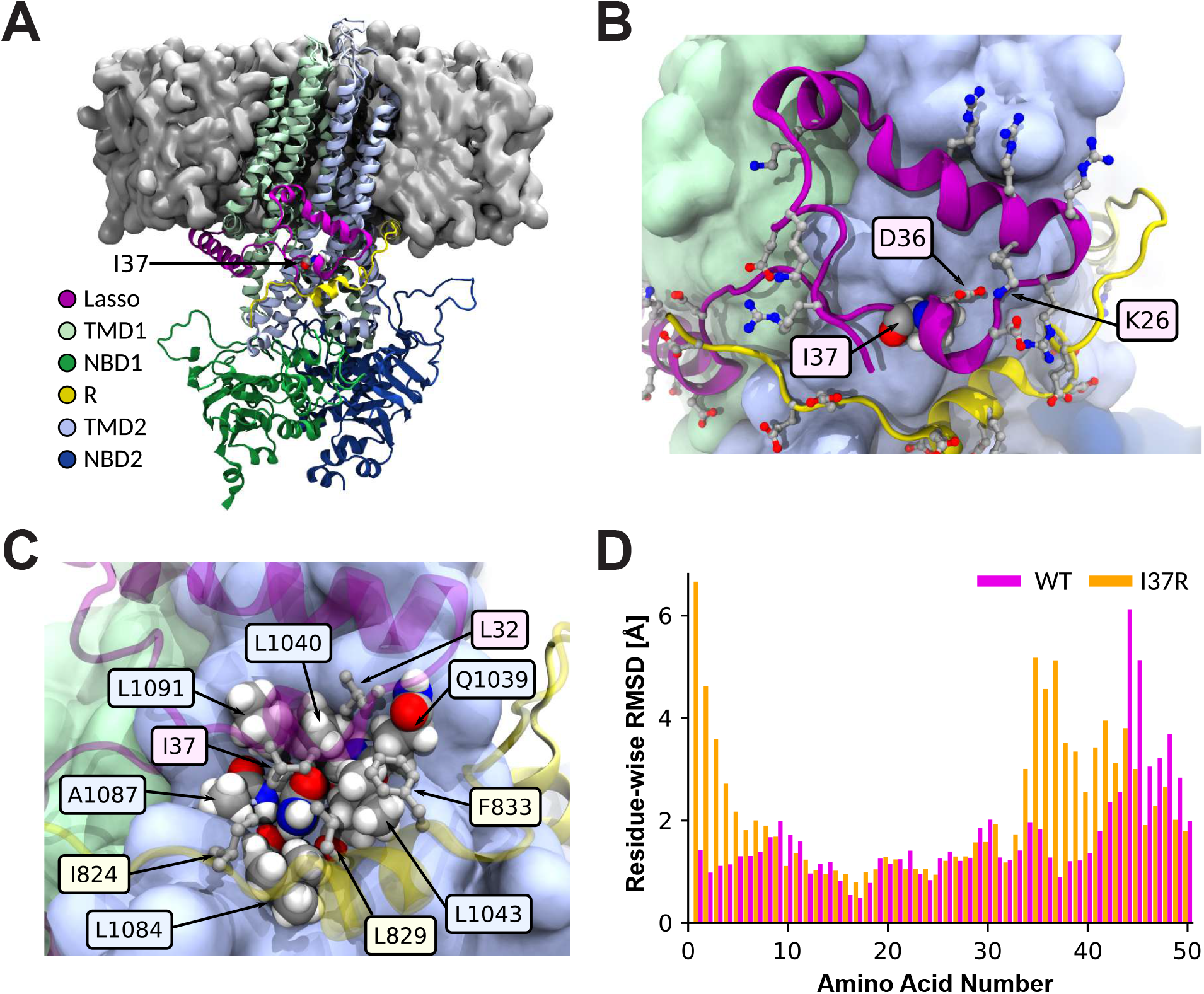
Placement of I37R within the lasso motif and the resulting changes to its conformation. (A) Ribbon structure of human CFTR, partially embedded within the plasma membrane (grey surface). I37 rendered as spheres. TMD: transmembrane domain; NBD: nucleotide binding domain; R: regulatory domain (R domain). (B) K26-D36 salt bridge stabilising the noose structure of the lasso motif. Charged amino acids depicted as balls and sticks. I37 rendered as spheres, and colour-coded by element (grey: C; white: H; red: O; blue: N). (C) I37 positioned within a hydrophobic pocket formed by amino acids from the lasso motif, TMD2, and poorly resolved R domain. Relevant amino acids labelled and depicted as spheres in TMD2, and as balls and sticks in lasso motif and R domain. (D) Residue-wise root-mean-square deviation (RMSD) to the C-alpha atoms of the WT-CFTR 6MSM model, measuring the conformational change of the WT (pink) and I37R mutant (orange) lasso after 2 µs simulations. Values are means sampled over the last µs of simulations.

Mutation of the evolutionarily conserved, non-polar and uncharged isoleucine (I) of I37 to a positively charged arginine (R) introduced an unstable lone charge into the hydrophobic pocket within the lasso motif noose. We hypothesised that this likely results in the rotation of the R37 side chain out of the hydrophobic pocket, and possible coordination with negative charges in the nearby R domain.

To identify a reasonable conformation of the mutant lasso motif, the WT 6MSM model was mutated to R37 and three 2 µs simulations were performed at physiological temperature (310 K). The R37 side chain rotated out of the hydrophobic pocket in only one of the three simulations. The difference between the root-mean-square deviation (RMSD) of the noose structure of I37R-CFTR compared to the WT was on average 2.8 Å at the amino acids M1-L6, and 1.8 Å at L34-S50 (**Fig 4D**). To confirm this observation, repeat simulations were performed at 350 K (40 degrees above physiological temperature), a temperature shown to accelerate the potential conformational transitions of proteins (Beckerman 2015). In these higher temperature simulations, the root-mean-square fluctuation (RMSF) of the region around amino acid 37 doubled in two out of three simulations, compared to WT-CFTR at 310K (**Supp material 8**). This confirmed the destabilisation of the lasso motif by I37R-CFTR. All WT-CFTR domains and the surrounding bilayer remained stable at the elevated temperature (**Supp material 9-10**).

### I37R mutation strengthens lasso motif interaction with the R domain

In the 6MSM structure, the R domain is largely unresolved with two exceptions: the first (Q637) and last (T845) amino acids that adjoin neighbouring domains, and the backbone atoms of a 17 amino acid segment. This latter segment consists of an eight amino acid disordered coil followed by a nine amino acid alpha-helix (Zhang, Liu, and Chen 2018). The alpha-helix was separated by approximately 10 Å (1 nm) minimum C-alpha distance to I37 in the lasso motif. This suggested a likely interaction between this segment of the R domain and I37, which necessitated partial modelling of the R domain (**Fig 5A**).

**Figure 5.**
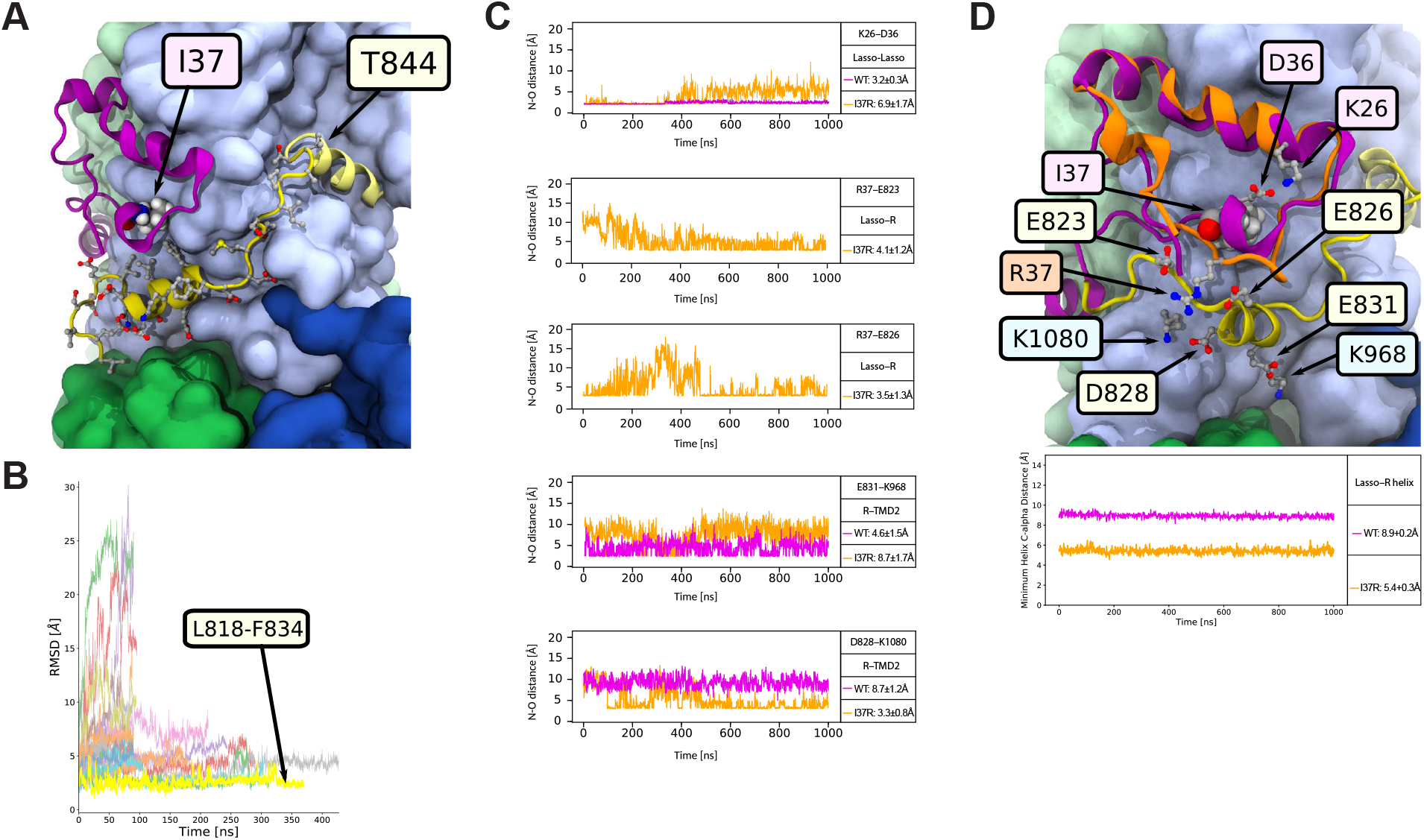
I37R interacts with a previously unresolved section of the R domain. (A) The reconstructed R domain amino acids (yellow), depicting the assignment of L818-F834 to the 17 amino acids with only the backbone resolved in the 6MSM structure, and the linking residues to T845 in TMD2. Lasso motif in purple. Side chains depicted as balls and sticks. (B) The stabilities of all 24 modelled R domain assignments, quantified by RMSD to the 6MSM structure. The most stable alignment of the 17 unidentified amino acids, L818-F834, is highlighted yellow. The full list of tested assignments is shown in Supplementary material 9. (C) The minimum N-O distance between newly formed and disrupted salt bridges in the I37R mutant. Distance less than 4 Å indicates direct contact. Values are means ± standard deviations (SD), sampled over the last 500 ns of simulations. (D) Conformational changes in the I37R mutant (orange) compared to WT (purple) lasso motif, which brings it closer to the R domain (yellow). Minimum C-alpha atom distance between amino acid 37 and the R domain helix (E826-F834) in I37R and WT.

Modelling of these 17 unidentified amino acids was performed by creating 24 different *in silico* models of this segment based on the 6MSM structure. In each model a unique 17 amino acid sequence was determined with a sliding window of one amino acid, starting backwards from amino acid T842 due to the alpha-helix’s 20 Å proximity to T845. The 17 amino acids were then connected to T845 with the missing linking amino acids. The structural stability of all 24 modelled segments was tested by performing up to 300 ns simulations for each model and comparing the backbone RMSD measurements against 6MSM (**Fig 5B, Supp material 11**). The model with the lowest RMSD (3 Å) and thus the highest stability was attained when L818-F834 was assigned to the unidentified 17 amino acids, of which the alpha-helix maps to E826-F834 (**Fig 5B, Supp material 11**). This assignment was corroborated by NMR measurements of the isolated R domain in solution, where the same segment retained partial helicity (Baker et al. 2007). Predictions of the structure of human CFTR by Alphafold2 also aligned with this assignment of primary structure to the unidentified amino acids (**Supp material 12**) (Jumper et al.). Several favourable interactions between this R domain model and other parts of the CFTR protein further supported this assignment (**Fig 4C, 5D**). Two hydrophobic amino acids (L829 and F833) contributed to the hydrophobic pocket that stabilised the lasso motif around I37. The negatively charged E831 formed a salt bridge with positively charged K968 in TMD2. Together, these interactions secured the R domain alpha-helix into position throughout an extended 2 μs simulation, resulting in a smaller minimum C-alpha distance to the lasso motif of 8.9±0.2 Å compared to the 10 Å in the 6MSM cryo-EM structure.

The reoriented R in position 37 in the I37R mutant protein, which pointed out of the hydrophobic pocket, rearranged the salt bridge network supporting the lasso motif by breaking the evolutionarily conserved salt bridge K26–D36. Two new salt bridges were formed, one with the negatively charged E823 and another with E826 of the R domain (**Fig 5C**). Furthermore, the E831–K968 salt bridge between the R and TMD2 domains in the WT was exchanged for a D828–K1080 salt bridge in I37R-CFTR (**Fig 5C**). The backbone motions required to accommodate these new charge interactions also perturbed parts of the lasso motif (**Supp material 8**) and R domain. The lasso N-terminus shifted its position towards the R domain and reduced the minimum C-alpha distance between them by 3.5 Å (**Fig 5D**). The overall result was a tighter coupling between the lasso and the R domain which is anticipated to inhibit the R domain movements required for channel gating.

## Discussion

We have described the functional and structural defects of I37R, a novel CF-causing mutation in the segment of the CFTR lasso motif which interacts with the R domain. These were compared to reference *CFTR* mutations which have known functional defects, either a CFTR folding/maturation (F508del/F508del) or a gating (G551D/F508del) defect. First, ICM performed in I37R/F508del rectal biopsies identified I37R confers high residual activity (50% of WT-CFTR activity). High baseline CFTR activity was similarly observed in FIS of I37R/F508del intestinal organoids and I_sc_ measurements in organoid-derived monolayers. Given we and others showed that F508del is a severe mutation which contributes little functional CFTR (Van Goor et al. 2011), this suggests that I37R mutation produces CFTR protein which localises to the epithelial cell surface. These observations are consistent with the patient’s mild CF clinical phenotypes (pancreatic sufficient with faecal elastase>500 μg/g, FEV_1_ z-score -0.11, 99% predicted).

We also characterised the response of I37R-CFTR to modulators (potentiators and correctors) in I37R/F508del intestinal organoids and organoid-derived monolayers. I37R was responsive to potentiators which improve CFTR gating function and a newly approved corrector (VX-445). Amongst the three potentiator agents tested, the response to VX-770 was minimal. The reason for the lack of efficacy of VX-770 is not known, since molecular modelling studies propose that VX-770 shares the same mechanism of action and binding sites with G1837 (Liu et al. 2019; Yeh et al. 2019). Both VX-770 and G1837 are proposed to potentiate CFTR by increasing channel open probability (Po) through stabilisation of the open-pore conformation, independent of NBD dimerization and ATP hydrolysis which normally controls channel gating (Yeh et al. 2017; Van Goor et al. 2009). However, the differing potentiator efficacies are not a new observation. G1837 was previously shown to be more potent and effective than VX-770 in human bronchial epithelial cells from a G551D/F508del and a R334W/F508del CF participant (Van der Plas et al. 2018; Gees et al. 2018). Similar observations were reported in heterologous HEK293 cells expressing Class III (G551D, G178R, S549N) and Class IV (R117H) CFTR mutants (Van der Plas et al. 2018; Gees et al. 2018). We conclude that perhaps G1837 has additional binding sites or actions distinct from VX-770, which in the case of I37R-CFTR, results in significant potentiation of the CFTR channel.

We further showed that dual potentiator combinations exerted synergistic restoration of CFTR activity in I37R/F508del organoids. This synergistic restoration is not exclusive to I37R-CFTR, since similar findings have been reported for other *CFTR* mutations responsive to potentiators (Veit et al. 2019; Phuan et al. 2018; Phuan et al. 2019; Dekkers, Van Mourik, et al. 2016). Synergism is commonly achieved when potentiators have distinct binding sites and mechanisms of actions. One potentiator could induce allosteric interactions that favour the activity of the other potentiator (Nussinov and Tsai 2013). The potentiator synergy observed in our dual potentiator combinations supports our hypothesis that G1837 may have additional binding sites or mechanisms of action to VX-770. While VX-770 has been shown to provide clinical benefit to patients with responsive mutations (McKone et al. 2014; Volkova et al. 2020; Berkers et al. 2020), it does not restore the Po of gating defect mutants (G551D-CFTR) to full WT-CFTR activity (Van Goor et al. 2009). This opens the possibility that using another potentiator with a different mechanism of action could complement VX-770 activity and increase CFTR activity beyond that of VX-770 monotherapy. While VX-770 and G1837 act independently of NBD dimerization and ATP hydrolysis (Yeh et al. 2017; Van Goor et al. 2009), genistein promotes ATP-dependent gating of CFTR by binding to the NBD1/2 interface and inhibiting ATP hydrolysis (Sohma, Yu, and Hwang 2013). Genistein has been demonstrated to increase VX-770-potentiated CFTR activity in intestinal organoids, even when VX-770 was used at near-saturating concentrations (Dekkers, Van Mourik, et al. 2016). Our observations reiterate and expand on these findings to suggest that potentiators with different mechanisms of action could provide synergistic restoration of CFTR activity to responsive *CFTR* mutations compared to potentiator monotherapy.

Chronic treatment with type III corrector VX-445 rescued CFTR activity in I37R/F508del organoids, while neither type I correctors (VX-809 or VX-661) rescued activity. This response is attributed to the I37R and not the F508del mutation in the I37R/F508del organoids, because VX-445 did not restore CFTR activity in F508del/F508del organoids. While VX-445 has been shown to have partial potentiator activity (Shaughnessy, Zeitlin, and Bratcher 2021; Laselva et al. 2021; Veit, Vaccarin, and Lukacs 2021), VX-445 did not potentiate CFTR activity in I37R/F508del organoids when administered acutely. This is the first study to interrogate the potentiator action of VX-445 in intestinal organoids, however previous studies have been performed in donor-derived bronchial and nasal epithelial cells, and immortalised cell lines. The higher correction efficacy of VX-445 when compared to VX-809/VX-661 has previously been shown, although this is likely to be dependent on the CFTR variant (Keating et al. 2018; Veit et al. 2021; Veit et al. 2020). For instance, direct binding of VX-445 to NBD1 to stabilise and prevent the domain unfolding may make it more effective in correcting CFTR mutations that impact NBD1 function (such as F508del located in NBD1).

The lack of I37R-CFTR correction by VX-809 or VX-661 could be attributed to the dependency of these modulators binding to and stabilising the TMD1. TMD1 function is modulated by interaction with lasso helix 2 (Lh2, aa A46–L61) as deletion of Lh2 from the WT CFTR was shown to completely abrogate VX-809-mediated CFTR maturation (Sabusap et al. 2021). MD studies showed that VX-809 occupancy at the TMD1 binding site causes the Lh2 to move, such that the network of salt bridges in Lh2 hold TMD1 (CL1) and TMD2 (CL4) in the correct orientation (Baatallah et al. 2021; Okiyoneda et al. 2013). This then allows for allosteric coupling between NBD1 and TMD1 or 2, which is important for cooperative domain folding of CFTR. In support of this, mutation of critical amino acids at the binding pocket of VX-809 on CFTR, or those involved in the architecture of this site, were shown to diminish the sensitivity to VX-809 correction. L53V and F87L mutations, which are located in the vicinity of the VX-809 binding site in the TMD1, were shown to prevent VX-809 correction in F508del HEK283 cells (Baatallah et al. 2021). Considering the above and since I37 is only a few amino acids away from the Lh2, it is plausible that the local conformational changes associated with the I37R mutation which we have identified in our study (**Fig 4D**) may disrupt the allosteric coupling between NBD1 and TMD1 or 2, preventing correction with type I correctors.

*CFTR* missense mutations in the lasso motif are not well characterised. This is because most of these mutations are rare, with an allele frequency of less than 0.01% in the CF population (**Supp material 1**). The only characterised missense mutations in the region of the lasso motif where I37 resides – between Lh1 (amino acid 19-29) and Lh2 (amino acid 46-61) – are R31C and R31L (CFTR2 2021; Jurkuvenaite et al. 2006). Experimental studies in heterologous COS-7 cells showed both mutations cause a mild processing defect and accelerated CFTR internalisation. Individuals heterozygous for these *CFTR* mutations are reported to have a mild disease phenotype with pancreatic sufficiency (Jurkuvenaite et al. 2006). One individual with the R31C/F508del *CFTR* genotype was reported to have a normal sweat chloride level (25 mmol/L) and nasal potential difference (Werlin et al. 2015). CFTR2 classifies R31C as a non-CF disease causing mutation. Notably, mild disease phenotypes (mild pulmonary symptoms, pancreatic sufficiency) are reported for several other lasso motif missense mutations including P5L, E56K and P67L (**Supp material 1**), as was found for the I37R/F508del participant in this study. This suggests that perhaps lasso motif mutations do not significantly impact the overall CFTR structure and function given its short length (69 of 1480 amino acids, 4.7%). It is also plausible that the role of the lasso motif could be compensated for by other CFTR domains.

To better understand the functional defect of I37R-CFTR, we used MD simulations to model the structural features of I37R and how they are altered relative to WT-CFTR. The amino acids 34-39 were shown to interact with the R domain in the phosphorylated, ATP-bound CFTR structure (Zhang, Liu, and Chen 2018). This interaction was absent in the closed conformation of CFTR (Zhang and Chen 2016), suggesting that the short region of amino acids 34-39 interacts with the R domain to regulate CFTR channel gating. We found that the disruption of the evolutionarily conserved K26-D36 salt bridge in I37R-CFTR brings the lasso motif closer to the R domain. We also found that the I37R side chain rotates out of its hydrophobic pocket to form interactions with negatively charged E823 and E826 on the R domain. We speculate that R37 clamps the lasso motif to the R domain, preventing the dynamic movement of the two domains necessary for a normal CFTR opening and closing cycle, thus causing a gating defect. This supports our functional observations, wherein I37R-CFTR demonstrated significant responsiveness to potentiator agents which are known to increase channel opening time. Furthermore, in the I37R-CFTR model, conformational changes in the lasso motif were also evident but were limited to short regions (M1-L6, L34-S50), indicating that the overall architecture of the CFTR protein remains largely intact. Additionally, our simulations did not show any change to the pore architecture of CFTR (**Supp material 13**).

The simulated structure in this work is of CFTR in its active state (Zhang, Liu, and Chen 2018). Because of this, we believe the pathogenic interactions discovered in this study have a significant contribution to the deleterious effects of the I37R mutation. However, the enhanced lasso motif-R domain interactions should be interpreted in the context of the μs timescales reachable by unbiased simulations. The lasso domain is known to exhibit conformational flexibility during both folding and functional stages of CFTR (Kleizen et al. 2021), which take place on timescales longer than is currently feasible to study in atomistic simulations. Therefore, there may be pathogenic interactions in I37R-CFTR in addition to the ones captured by the simulation of this particular CFTR structure.

The I37R/F508del participant in this study will only meet the Therapeutic Goods Administration (Australia) requirements for treatment with Trikafta/Kaftrio^®^triple combination therapy once he turns 12 years old given the single copy of the F508del mutation. He is not eligible for single potentiator therapy or corrector/potentiator combinations of lumacaftor/ivacaftor or tezacaftor/ivacaftor. This emphasises the importance of characterising the structural and functional defects of ultra-rare *CFTR* mutations together with the assessment of *in vitro* response to modulator drugs in patient-derived cell models to build the case for access to treatment with available modulators through precision medicine health technology assessment pathways. Furthermore, when multiple CFTR modulators are available to CF patients, determining the best modulator for patients with a rare mutation not investigated in a clinical trial may be supported using *in vitro* personalised cell models.

## Author contributions

Conception and design: SAW, AJ. Recruitment and consent: LF, SAW. Collection of rectal biopsies: CYO, LF. Ion transport assay: NTA. Culturing of organoids: NTA, SLW, SAW. FIS microscopy: IS, KA, SLW. FIS scripts: MC, RW. FIS analysis: NTA, SLW. Immunofluorescence microscopy: SLW. Western blot: SLW. Molecular Dynamics: MA, PC, RG, SK. CFTR sequence alignment: AC. Figure preparation: SLW, MA, NTA, AC, KA, SAW. Writing – original draft: SLW, MA, SAW. Review and editing: SAW, KA, RG, SK, LF with intellectual input from all other authors. Supervision: SAW, SK.

## Acknowledgements

We thank the study participants and their families for their contributions. We also thank Sydney Children’s Hospital’s (SCH) Randwick respiratory department especially Leanne Plush, Amanda Thompson, Roxanne Strachan and Rhonda Bell for the organisation and collection of participant biospecimens. SAW is supported by an Australian National Health and Medical Research Council grant NHMRC_APP1188987. MA acknowledges support of a top-up scholarship from Cystic Fibrosis Australia. MA and KA are supported by Australian Government Research Training Program Scholarship. We acknowledge the Hubrecht Institute for the generous provision of the L-Wnt3A cell line. Computations were performed on the Gadi HPC at the National Computational Infrastructure Centre in Canberra and Artemis at the Sydney Informatics Hub in The University of Sydney.

## STAR Methods

### EXPERIMENTAL MODEL AND SUBJECT DETAILS

#### Participants biospecimen collection

Rectal biopsies were collected from CF participants with I37R/F508del (n=1), F508del/F508del (n=6) and G551D/F508del (n=2) *CFTR* genotypes, as well as WT-*CFTR* control participants (n=3) (**Supplementary material 2**). Samples were taken during investigative or surveillance endoscopy. This study was approved by the Sydney Children’s Hospital Ethics Review Board (HREC/16/SCHN/120). Written informed consent was obtained from all participating subjects or their legal guardians.

#### Intestinal organoid culture from rectal biopsies

Organoid cultures were established from crypts isolated from four to six rectal biopsies as described previously (Dekkers, Berkers, et al. 2016). Briefly, rectal biopsies were washed with cold PBS, and incubated with 10 mM EDTA (Life Technologies 15575-020) in PBS at 4°C for 120 min on a tube rotator. EDTA solution was discarded and crypts were dislodged by vigorous pipetting of biopsies in cold PBS. Isolated crypts were seeded in 70% matrigel (Growth factor reduced, phenol-free; Corning 356231) in 24-well plates at a density of ∼10 – 30 crypts in 3×10 µl matrigel droplets per well. Media change was performed every second day and organoids were passaged 1:3 after 7 – 10 days of culture.

#### Organoid-derived monolayers cultures

Monolayers cultures were created from intestinal organoids as described previously (Zomer-van Ommen et al. 2018). Briefly, 7-day-old organoids were dissociated into single cells using TrypLE Express Enzyme (Life Technologies 12605-010) at 37°C for 2×2 min. Mechanical disruption was performed after each incubation period. Cells were then seeded on collagen I-coated (Advanced Biomatrix 5005) Transwell 6.5mm, 0.4 µm pore polyester membrane inserts (Sigma CLS3470) at a density of 250,000 cells per insert. Cells were cultured with organoid culture media supplemented with 10 µM Y-27632 for 48 h. Media change (without Y-27632) was performed every second day for 7 days.

### METHOD DETAILS

#### Intestinal current measurement

Superficial rectal mucosa samples (2 – 4 per donor) were freshly obtained using biopsy forceps (CK Surgitech NBF53-11023230) and placed in cold RPMI1640 media (Sigma R5886) with 5% FBS. Intestinal current measurements were performed under voltage-clamp conditions using VCC MC8 Ussing chambers (Physiologic Instruments, San Diego, CA) (De Jonge et al. 2004; Li, Sheppard, and Hug 2004; Derichs et al. 2010). Biopsy tissues were bathed in Ringer solution containing (mM): 145□NaCl, 3.3□K_2_HPO_4_, 0.4 KH_2_PO_4_, 10 D-Glucose, 10 NaHCO_3_, 1.2 MgCl_2_ and 1.2□CaCl_2_. Ringer solutions were continuously gassed with 95% O_2_-5% CO_2_ and maintained at 37°C. 10 µM indomethacin was added to both apical and basal chambers, and tissues were stabilised for 40 min. Tissues were then treated with pharmacological compounds (in order): 100 µM amiloride (apical) to inhibit epithelial sodium channel (ENaC)-mediated Na^+^ flux, 10 µM forskolin + 100 µM IBMX cocktail (apical and basal) to induce cAMP activation of CFTR, 100 µM carbachol (basal) to increase intracellular Ca^2+^ levels and activate basolateral Ca^2+^-dependent K^+^ channels and 100 µM bumetanide (basal) to inhibit basolateral Na^+^/K^+^/2Cl^-^ (NKCC) co-transporter.

#### Forskolin-induced swelling assay

Measurement of fsk-induced swelling (FIS) assay was performed as described previously (Wong et al. 2021). Passage 3-15 organoids were seeded in 96-well plates, in 4 µl 70% matrigel droplet per well containing ∼25–30 organoids. The next day, organoids were incubated with 1.84 µM calcein green (Thermo Fisher Scientific C3100MP) for at least 30 min prior to addition of fsk at 0.02, 0.128, 0.8 or 5 µM concentrations, to determine cell viability. For CFTR potentiation, a single potentiator (3 µM VX-770 or 3 µM G1837 or 50 µM Gen) or dual potentiators (VX-770+G1837 or VX-770+Gen or G1837+Gen) was added together with fsk. Time-lapse images of organoid swelling were acquired at 10-min intervals for 60 min at 37°C using Zeiss Axio Observer Z.1 inverted microscope (Carl Zeiss, Jena, Germany), on an EC Plan-Neofluar 5x/0.16 M27 dry objective. Organoids were pre-incubated with 3 µM VX-809 or 3 µM VX-661 or 3 µM VX-445 or 3 µM VX-445+18 µM VX-661 for 24 h prior to FIS for CFTR correction where indicated. Three wells were used per condition and each participant’s FIS experiment was repeated 3 to 4 times.

#### Quantification of forskolin-induced swelling

Organoid swelling was quantified using a custom-built script as described previously (Wong et al. 2021). A segmentation strategy implemented using ImageJ/Fiji was performed on brightfield images. The raw image was processed with a gaussian blur (s=1.3) to reduce noise. After the directionality and magnitude of the local gradient was identified, pixels were classified as either ‘Background’, ‘Ridge’, ‘Valley’, ‘Rising’ or ‘Falling’ dependent on their neighbouring pixels along the previously calculated local directionality. Clean-up filters were applied that remove noise and small objects, such as ridges that only touched background pixels, and erosions to decrease rising and falling edges to better approximate object boundaries (‘Peaks’). A size exclusion was applied that would discriminate debris in the sample preparation from organoids of interest. This segmentation strategy was used to identify area covered by organoid at each time point. The total surface area of organoid at 10-min intervals over 60 min post-fsk stimulation were calculated and normalized against t=0 to render the relative amount of swelling from t=0. The area under the curve, AUC (calculated increase in organoid surface area from t=0 to t=60; baseline=100%) was then calculated using GraphPad Prism software.

#### Quantification of CFTR-mediated ion transport in organoid-derived monolayers

Short circuit current (I_sc_) measurements were performed under voltage-clamp conditions using VCC MC8 Ussing chambers (Physiologic Instruments, San Diego, CA). Cells were bathed in 20□mM HEPES buffered-Ringer solution containing (mM): 120□NaCl, 0.8□K_2_HPO_4_, 5 D-Glucose, 1.2 MgCl_2_ and 1.2□CaCl_2_. Ringer solutions were continuously gassed with 95% O_2_-5% CO_2_ and maintained at 37°C. 10 µM indomethacin was added to both apical and basal chambers and cells were stabilised for 15 min. Cells were then treated with pharmacological compounds (in order): 100 µM amiloride (apical) to inhibit epithelial sodium channel (ENaC)-mediated Na^+^ flux, vehicle control 0.01% DMSO or 10 µM G1837 (apical) to potentiate cAMP-activated currents, 5 µM forskolin (basal) to induce cAMP activation of CFTR, 30 µM CFTR_inh_-172 (apical) to inhibit CFTR-specific currents and 100 µM ATP (apical) to activate calcium-activated chloride currents. I_sc_ in response to forskolin was considered as baseline activity (ΔI_sc-Fsk_) and I_sc_ in response to forskolin and potentiator (ΔI_sc-Fsk+Pot_) was used as the measure of modulator response.

#### Immunofluorescence

A rectal biopsy from a I37R/F508del participant was embedded in Tissue-Tek Optimal Cutting Temperature (OCT) compound (Sakura Finetek, CA) and snap frozen prior to storage at -80° C. The frozen biopsy was cut into 4 μm slice sections, and the sections were fixed in ice-cold methanol for 15 min. Intestinal organoids cultured from the I37R/F508del participant and organoid-derived monolayers cultured from a F508del/F508del participant were fixed in 4% paraformaldehyde and ice-cold methanol respectively for 15 min. Fixed samples were blocked using IF buffer (0.1% BSA, 0.2% Triton and 0.05% Tween 20 in PBS) with 10% normal goat serum (Sigma G9023) for 1 h at room temperature before incubation in primary antibodies overnight at 4°C. The biopsy section was stained with CFTR (1:50, Abcam ab2784) and E-cadherin (1:100, Cell Signalling 3195) antibodies. Intestinal organoids were stained with Ki67 (1:250, Abcam ab15580) and E-cadherin (1:250, Life Technologies 13-1700) antibodies. Organoid-derived monolayers were stained with ZO-1 (1:250, Life Technologies 61-7300) and E-cadherin (1:250, Life Technologies 13-1700) antibodies. On the following day, samples were washed with IF buffer 3 times, 5 min each and incubated with Alexa Fluor conjugated secondary antibodies (1:500, Life Technologies A-11029, A-21329) for 1 h at room temperature. Samples were mounted with Vectashield hardset antifade mounting medium containing DAPI (Vector Laboratories H-1500). Images were acquired using Leica TCS SP8 DLS confocal microscope (Leica Microsystems, Wetzlar, Germany), either on a 63x/1.4 or a 20x/0.75 objective. Images were processed using ImageJ (National Institutes of Health, Bethesda, MD).

#### Western blotting

Intestinal organoids were lysed with TNI lysis buffer (0.5% Igepal CA-630, 50 mM Tris pH 7.5, 250 mM NaCl, 1 mM EDTA) (Pankow et al. 2015) containing protease inhibitor cocktail (Roche 04693159001) on ice for 30 min. Lysates were then sonicated using the Bioruptor Pico (Diagenode, Liège, Belgium) at 4°C for 20 cycles of 30 sec on and 30 sec off. Lysates were spun down at 14,000 rpm at 4°C for 20 min and protein concentrations were determined using the BCA Protein Assay Kit (Thermo Fisher Scientific 23225). Lysates (100 µg per sample) were separated using NuPAGE 3 – 8% Tris-Acetate gels (Thermo Fisher Scientific EA0375BOX) at 100 V for 30 min, followed by 150 V until separation was complete. Proteins were transferred onto a nitrocellulose membrane using wet transfer at 20 V for 1 h at RT. The membrane was then incubated in 5% non-fat dry milk in phosphate-buffered saline containing 0.1% Tween (PBST) for 1 h at RT. CFTR bands were detected using anti-CFTR antibody 596 (1:500; University of North Carolina, Chapel Hill and Cystic Fibrosis Foundation) incubated at 4°C overnight. Protein bands were visualised using ECL Select detection reagent (Cytiva RPN2235) on the ImageQuant LAS 4000 (GE Healthcare, Chicago, IL). Calnexin was used as the loading control, detected using anti-calnexin antibody (1:1000; Cell Signalling Technology 2679). Protein band densitometry was performed using ImageJ (National Institutes of Health, Bethesda, MD). CFTR maturation in I37R/F508del and F508del/F508del organoids were estimated by measuring the level of mature mutant CFTR (band C) as a percentage of mature CFTR from WT organoids (% normal CFTR) (Van Goor et al. 2014).

### COMPUTATIONAL METHOD DETAILS

#### *In silico* system composition

A 1-palmitoyl-2-oleoyl-sn-glycero-3-phosphocholine (POPC) bilayer was generated using the VMD membrane builder plugin (Humphrey, Dalke, and Schulten 1996) in which a model based on the phosphorylated human CFTR channel (PDB ID: 6MSM) was embedded (Zhang, Liu, and Chen 2018). The system was solvated with TIP3P water and neutralised with 0.15 M of potassium chloride ions (Mark and Nilsson 2001). The WT-CFTR system included 236 POPC molecules, 128 potassium ions, 140 chloride ions and 44503 water molecules.

#### Extended 6MSM structure: modelling the unidentified section of the R domain

The 6MSM structure was extended in order to resolve a previously unassigned section in the R domain. The R domain is 227 residues long (F630-H856) and is largely disordered (Bozoky et al. 2013). In the 6MSM structure, the sidechains of 17 residues of this domain are labelled “UNKNOWN”, due to inadequate electron density in the region. The first 8 residues are unstructured while the next 9 residues form an alpha helix. The distance between the end of the helix and the first visible residue in TMD2 (T845) is 20 Å (Zhang, Liu, and Chen 2018). Using VMD’s autopsf plugin (Humphrey, Dalke, and Schulten 1996) we populated the side chains of the unknown section. Modeller 9.19 was then used to link the R domain to TMD2 at T845 (Sali and Blundell 1993). 24 possible primary structure alignments of this region were simulated. The Root Mean Squared Deviation (RMSD) of the backbone alpha carbon atoms of the extended section with respect to the 6MSM structure was calculated over 300 ns of MD simulations. The most stable alignment was chosen from the lowest RMSD compared to the 6MSM structure. The most stable configuration was capped with the neutral forms of the C and N termini and incorporated into our CFTR model. Four other missing loops namely residues 410-434, 890-899, 1174-1201, 1452-1480 were reconstructed using Modeller 9.19, based on visual analysis and the lowest discrete optimised protein energy (DOPE) score (Shen and Sali 2006). The N and C termini of the CFTR model were capped with the physiological, charged termini.

#### Molecular dynamics simulation protocols

The 6MSM structure carries an engineered mutation to avoid the hydrolysis of the bound ATP, giving it a longer lifetime in the open conformation (E1371Q). This mutation was corrected to match the WT-CFTR sequence using the mutator plugin of VMD. The I37R missense mutation was constructed in the same way. Gromacs v2019.3 with the CHARMM36m forcefield was used for all MD simulations (Abraham et al. 2015; Huang et al. 2017). Minimisation via a steepest descent algorithm was performed until all forces were below 24 kcal/mol/Å. This was followed by relaxation simulations of all heavy atoms in the system starting with a restraint of 10 kcal/mol/Å^2^ and then halving this restraint every 200 ps in 15 iterations. Relaxation and production were run with 1 and 2 fs time steps, respectively. Relaxation was followed by 5 ns of equilibration. During relaxation, a Berendsen thermostat and barostat were applied, and for production a Nosé-Hoover and Parrinello-Rahman thermostat and barostat were applied respectively (Parrinello and Rahman 1981; Nosé and Klein 1983; Berendsen et al. 1984). To maintain the area per lipid (APL) properties of the POPC membrane at experimental values during production runs, pressure coupling was applied in the z-direction normal to the membrane bilayer while the x-y dimensions of the cubic simulation volume was fixed (Klauda et al. 2010). While semi-isotropic pressure coupling better replicates membrane environments (Pandit and Scott 2008), this constant area approach was adopted to circumvent an issue with GROMACS 2019.3reb (https://gitlab.com/gromacs/gromacs/-/issues/2867). Production runs were extended up to 2 μs at 310 K with three replicates for all simple MD simulations. The last 1 μs of the longest simulations for each system were selected for further analysis. This was the longest time feasible to simulate with available computational resources. All RMSDs were calculated using the positions of alpha carbons with reference to the 6MSM experimental structure (Zhang, Liu, and Chen 2018). Analysis scripts were written in python using the MDAnalysis library (Michaud-Agrawal et al. 2011; Gowers et al. 2016). Bilayer thickness and area per lipid were calculated with the FATSLiM software package (Buchoux 2017).

#### Mathematical Formulae

Root Mean Square Deviation (RMSD)

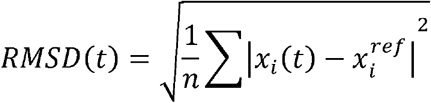

Root Mean Square Fluctuation (RMSF)

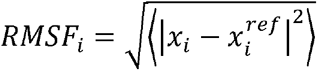

Angular brackets indicate a time average.

## QUANTIFICATION AND STATISTICAL ANALYSIS

Data for Fig 1A, G and Fig 2D are presented as dot plots with mean ± standard error of the mean (SEM). Data for Fig 1D, Fig 2A and E are presented as line graphs with mean ± standard deviation (SD). Data in Fig 1E, Fig 2C, G, Fig 3A and B are presented as violin plots with mean. One-way analysis of variance (ANOVA) was used to determine statistical differences. Statistical analysis was performed with GraphPad Prism software v9.0.1. A *P*-value of less than 0.05 was considered to be statistically significant.

## DATA AND SOFTWARE AVAILABILITY

An equilibrated model of CFTR with the missing R domain fragment has been deposited in Zenodo (10.5281/zenodo.5642866) alongside the workflow to simulate and analyse it.

